# Live cell monitoring of double strand breaks in *S. cerevisiae*

**DOI:** 10.1101/265611

**Authors:** David P. Waterman, Cheng-Sheng Lee, Michael Tsabar, Felix Zhou, Vinay V. Eapen, Allison Mazzella, James E. Haber

**Affiliations:** Rosenstiel Basic Medical Science Center and the Department of Biology, Brandeis University, Waltham, MA, 02454

## Abstract

We have used two different live-cell fluorescent protein markers to monitor the formation and localization of double-strand breaks (DSBs) in budding yeast. Using GFP derivatives of the Rad51 recombination protein or the Ddc2 checkpoint protein, we find that cells with three site-specific DSBs, on different chromosomes, usually display 2 or 3 foci that coalesce and dissociate. Rad51-GFP, by itself, is unable to repair DSBs by homologous recombination in mitotic cells, but is able to form foci and allow repair when heterozygous with a wild type Rad51 protein. The kinetics of disappearance of Rad51-GFP foci parallels the completion of DSB repair. However, in meiosis, Rad51-GFP is proficient when homozygous. Using Ddc2-GFP, we conclude that co-localization of foci following 3 DSBs does not represent formation of a homologous recombination "repair center," as the same distribution of Ddc2-GFP foci was found in the presence or absence of the Rad52 protein. The maintenance of separate DSB foci and much of their dynamics depend on functional microtubules, as addition of nocodazole resulted in a greater population of cells displaying a single focus.

**Author Summary:** Double strand breaks (DSBs) pose the greatest threat to the fidelity of an organism’s genome. While much work has been done on the mechanisms of DSB repair, the arrangement and interaction of multiple DSBs within a single cell remain unclear. Using two live-cell fluorescent DSB markers, we show that cells with 3 site-specific DSBs usually form 2 or 3 foci what can coalesce into fewer foci but also dissociate. The aggregation of DSBs into a single focus does not depend on the Rad52 recombination protein, suggesting that there is no “repair center” for homologous recombination. DSB foci are highly dynamic and their dynamic nature is dependent on microtubules.

## Introduction

The process of repairing chromosomal double-strand breaks by Rad51-and Rad52-mediated homologous recombination in budding yeast has been defined by a combination of in vitro analysis of purified recombination proteins (1-3) and from “in vivo biochemistry” analyses of the kinetics of repair of site-specific DSBs (4). Cleaved DNA ends are attacked by several 5’ to 3’ exonucleases to produce long 3’-ended single-strand DNA (ssDNA) tails, which are initially coated by the single-strand binding complex, RPA (5, 6). RPA is displaced by Rad51 recombinase through the action of mediator proteins, including Rad52, creating a nucleoprotein filament composed primarily of Rad51 but also its paralogs, the Rad55-Rad57 heterodimer (7-9). The Rad51 filament engages in a genome-wide search for a homologous sequence that could be on a sister chromatid, a homologous chromosome or at an ectopic location. Once the donor sequence is encountered, Rad51 catalyzes strand exchange to form a D-loop intermediate, the initial step in repair. The 3’ end of the invading strand then acts as a primer to initiate new DNA synthesis that leads to repair of the DSB via several pathways including gene conversion via synthesis-dependent strand annealing or a double Holliday junction pathway. A combination of Southern blot, PCR and chromatin immunoprecipitation (ChIP) experiments have shown that DSB repair proceeds by a series of kinetically slow steps, taking more than an hour to complete (reviewed in (4)).

In haploid cells, successful recombination with an ectopic donor sequence is strongly dictated by the contact probability of sequences within different chromosomes (10,11). When the ends of the DSB fail to encounter a donor, or in the case where there is no donor, an unrepaired break eventually enters a different pathway, where it associates with the nuclear envelope through its association with the nuclear envelope protein Mps3 (12). Localization to the envelope may alter further end-resection and may facilitate joining of DSB ends by nonhomologous end-joining (13).

One approach to the study of DSB repair in budding yeast has been the use of live-cell microscopy to monitor the behavior of different fluorescently tagged repair-associated proteins. The most thoroughly studied is Rad52, the key mediator for the assembly of the Rad51 filament, but which is also critical in later strand-annealing steps(14). Strikingly when there are multiple DSBs, created by ionizing radiation or by site-specific endonucleases, there often appears to be a single fluorescent Rad52 focus. This observation has led to the idea that there could be a “repair center” where recombination proteins might accumulate to facilitate DSB repair (15). However, immunofluorescent staining of spread nuclei with multiple DSBs found that the number of foci directly correlated with the number of DSBs (16)

A limitation in extending these studies has been the absence of other live-cell markers to follow repair. To this end, we constructed and characterized a Rad51-GFP fusion protein. Previously, a Rad51-GFP fusion was characterized in *Arabidopsis,* where it proved to be defective in mitotic DSB repair, but competent in meiosis (17). This phenotype resembles the “site II” mutation of *Saccharomyces cerevisiae* Rad51, which can bind ssDNA but is unable to bind dsDNA and thus fails to complete strand invasion and DSB repair in mitotic cells (18,19). Similar results were obtained using a human isoform of Rad51-GFP *in vitro* (20). In fission yeast, scRad51’s homolog Rhp51 when fused with CFP proved to be UV sensitive and incapable of carrying our repair on its own, but this defect was complimented by expression of wild type Rhp51 (21). Here we show that yeast Rad51-GFP binds to site-specific DSBs in mitotic cells but cannot catalyze homologous recombination when it is the only allele present; however, it is not dominant-negative – as is a similar construct in *Arabidopsis* (17). Consequently, Rad51-GFP can be used to follow GFP-labeled filaments that are engaged in functional recombination. In meiosis, budding yeast Rad51 acts as an auxiliary factor with the Rad51 homolog, Dmc1, and the site II mutant is competent for meiotic recombination (18). As with the *Arabidopsis* construct, yeast Rad51-GFP is competent for meiosis. Thus, we have developed a live-cell reporter for Rad51 in response to DSBs in both mitotic and meiotic cells where recombination can be induced synchronously.

Using either Rad51-GFP or a GFP fusion of the DNA damage checkpoint protein Ddc2, yeast’s homolog of the ATRIP protein that has been previously shown to bind near a DSB and to recruit Mec1^ATR^ kinase (22, 23), we show that cells which have multiple site-specific DSBs form multiple, highly dynamic GFP foci that coalesce and separate. In the majority of these cases, there are also multiple Rad52 foci, although some limitation in Rad52-RFP expression or a propensity for self-aggregation appears to restrict the number of Rad52-RFP foci even when there are distinct GFP foci. These results suggest that multiple DSBs do not generally form a Rad52-dependent repair center.

## Results

### Rad51-GFP forms a DNA damage-dependent focus

An ideal tool for monitoring DSB formation and repair would be a fluorescent protein that performs a central role in homologous recombination. We created a Rad51-GFP fusion construct utilizing a-SSGSSG-linker, which we have previously used to increase the functionality of other fusion proteins (24). We integrated this construct at the C-terminus of the genomic copy of *RAD51* in the donorless, galactose inducible HO-inducible strain JKM179 in which a single irreparable DSB is induced upon addition of galactose (25). More than 70% of cells displayed a single GFP focus within 3 h after inducing HO expression, increasing to >90% by 5 h (Figure 1D and S1A). Rad51-GFP foci were absent in *rad52Δ* cells or in cells that lack an HO cleavage site (Figures 1 and S1). When Rad51-GFP was coexpressed with Rad52-RFP, green and red foci colocalized (Figure 1F, S1D).

Rad51 has been shown to increase in abundance after DNA damage (26, 27). Such an increase is evident comparing the total nuclear intensity of Rad51-GFP in cells with a DSB (with or without Rad52) compared to cells lacking the HO cleavage site (Figure 1E).

**Fig 1.**
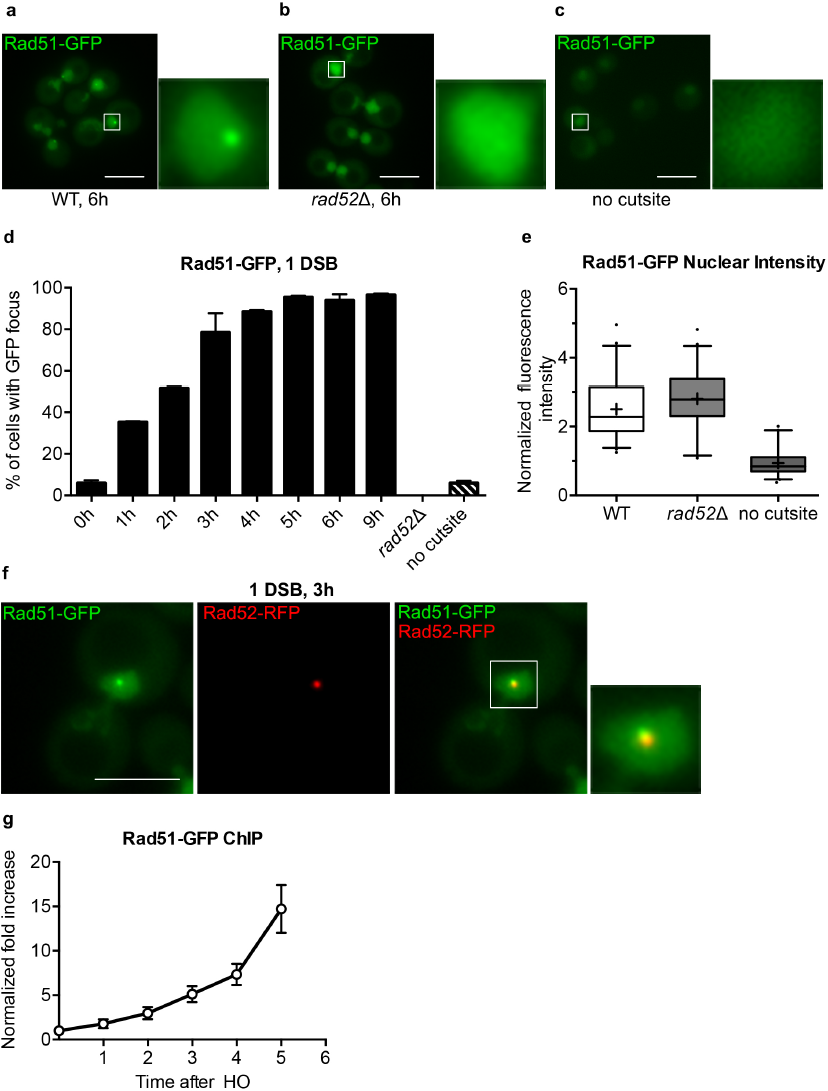
Rad51-GFP forms a DSB-dependent focus. **a)** Representative images of strain DW58 expressing endogenous Rad51-GFP 6 h after HO induction. Magnification of white boxed nucleus shown to the right. Scale bar = 5 µm. **b)** Representative images of strain DW88 (*rad52Δ*) prepared as in (a), **c)** Representative images of strain DW94 (no HO cut site) prepared as in (a), **d)** Quantification of the number of cells displaying Rad51-GFP foci in strain DW58, DW88, and DW94 atthe indicated time. **e)** Background-subtracted fluorescence intensities of nuclei in strains DW58 (WT), DW88 (*rad52Δ*), and DW94 (WT, no HO cut site) 6 h after HO induction as described in (a). Box plots display the median (black bar), mean (+), 25^th^ and 75^th^ percentiles (box ranges), 5th and 95^th^percentiles (whiskers), and outliers (dots). **f)** Representative image from strain DW89 expressing endogenous Rad51-GFP and Rad52-RFP from its endogenous Promoter on a low copy plasmid 3 h after HO induction prepared as in (a) **g)** Quantification of Rad51 ChIP signal atthe indicated time after induction of HO. Error bars represent the SD of three biological replicates of >150 cells per experiment.

To test directly if Rad51-GFP was bound to the DNA around the DSB, we performed chromatin immunoprecipitation using an antibody recognizing Rad51 to assay Rad51-GFP accumulation 5 kb from the DSB induced in the *MATα* locus in a derivative of strain JKM179, lacking donor sequences, as described previously (28). As shown in Figure 1G, Rad51-GFP binding 5 kb from the unrepaired DSB end increased steadily over 6 h. Rad51-GFP binding appears similar in its kinetics to wild type Rad51, as measured previously (19, 28). Therefore, Rad51-GFP effectively binds to resected DNA around a DSB and thus shows promise for further live cell studies.

### Rad51-GFP cannot repair DSBs by homologous recombination in mitotic cells, but it is not dominant negative

Next, we sought to determine whether Rad51-GFP was functional with regard to the DNA damage checkpoint and to DSB repair. Cells that suffer an irreparable DSB arrest for 9-12 h through activation of the DNA damage checkpoint. After about 12 h, and without repairing the DSB, yeast cells switch off the checkpoint and proceed through mitosis in a process called adaptation (25, 29, 30). Adaptation requires Rad51, but not Rad52; in *rad51*Δ most cells permanently arrest prior to mitosis after a DSB (31). However, Rad51-GFP cells are capable of adapting similar to wild type (Figure 2A).

**Fig 2.**
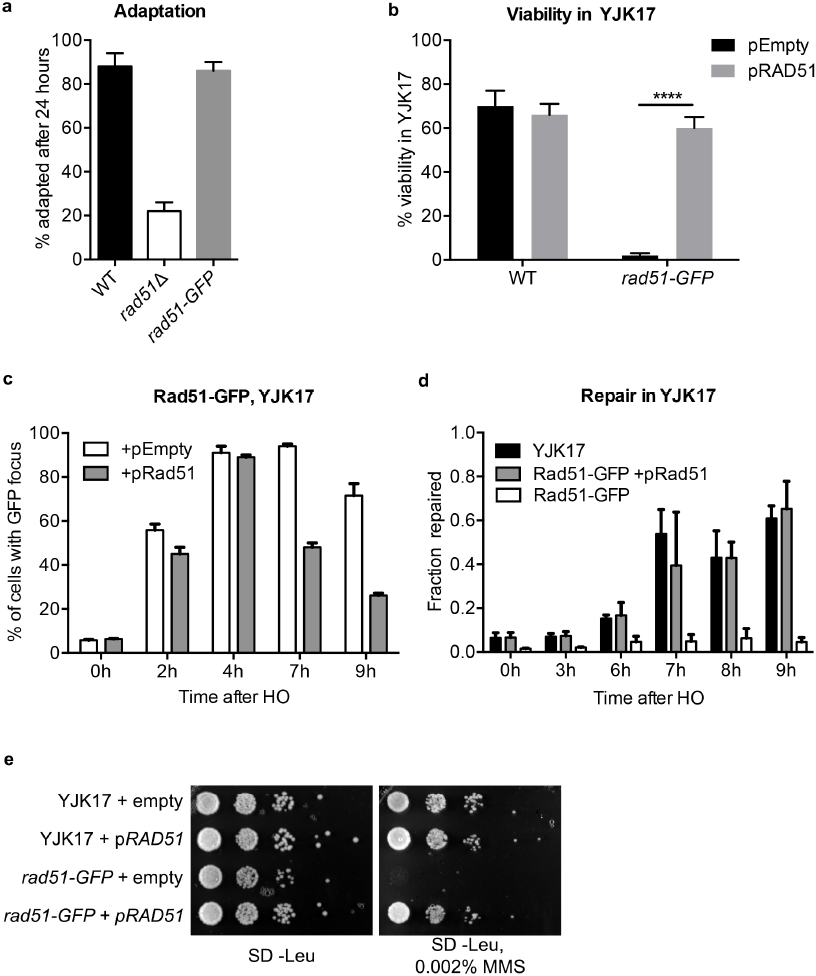
Rad51-GFP is not dominant negative. **a)** Quantification of the percentage of cells that adapt after 24 h after HO induction in the indicated strain. **b)** Quantification of the percent of viable cells following HO induction and repair through ectopic gene conversion in the indicated derivative of YJK17. Student’s t test **** p ≤ 0.0001.**c)** Quantification of the percentage of cells displaying a focus in the indicated derivative of YJK17 at the indicated time, **d)** qPCR analysis of the timing of DSB repair by gene conversion in the indicated derivatives of YJK17. **e)** Spot dilution assay without and with 0.002% MMS. Error bars represent the SD of three biological replicates.

In the assays described thus far, the DSBs were irreparable by HR because of the lack of a donor template. To investigate the ability of Rad51-GFP to participate in HR, we turned to strain YJK17, in which there is a DSB at *MATα* on Chr3 and a single ectopic *MAT***a**-inc donor sequence on Chr5 (32). An HO break is repaired in roughly 80% of cells over the course of 6-9 h. YJK17 carrying Rad51-GFP failed to repair the DSB (Fig. 2B). Given the multimeric nature of the Rad51 filament and that many Rad51 mutations are dominant-negative (33, 34) we asked if Rad51-GFP is dominant negative. We found that HO-induced DSB repair in YJK17 was repair-proficient after introducing wild type Rad51 on a centromeric plasmid, expressed from the its own promoter (Figure 2B). The kinetics of repair, monitored by qPCR, were very similar for Rad51-GFP complemented by *RAD51* compared to wild type (Figure 2D). In parallel with repair, the percent of cells displaying a GFP focus decreased from 80% at 4 h to ~50% by 7 h and fewer than 30% by 9 h, whereas without the complementing Rad51, foci persisted (Figure 2C). This decreased correlated with the timing of repair as monitored by qPCR (Figure 2C).

Further evidence that Rad51-GFP is not dominant-negative was found by monitoring cells exposed to 0.002% MMS, which was lethal to Rad51-GFP cells but not to wild type (Figure 2E). The sensitivity of the Rad51-GFP strain was rescued by providing wild type *RAD51,* expressed from its own promoter, on a centromere-containing plasmid. These data suggest that Rad51-GFP is capable of binding DNA even in the presence of wild type Rad51 and the GFP fusion’s loss of function can be complimented by expression of wild type Rad51.

### Rad51-GFP is competent in meiosis

*Arabidopsis* Rad51-GFP proved to be meiosis-competent even though it blocked mitotic recombination (17). As noted above, this phenotype resembles a Rad51 “site II” mutation in budding yeast (18). In meiosis, the critical functions of strand exchange depend on Rad51’s homolog, Dmc1, with Rad51 acting in an apparently allosteric fashion. We found that Rad51-GFP is meiosis-proficient. Diploids homozygous or heterozygous for Rad51-GFP produced the same percentage of as wild type. (Figure 3A). After tetrad dissection, spores resulting from diploids homozygous for Rad51-GFP exhibited a 40% reduction in spore viability, but nevertheless 60% of spores were viable (Figure 3B). Thus, *S. cerevisiae* Rad51-GFP strongly resembles a site II mutation (18).

**Fig 3.**
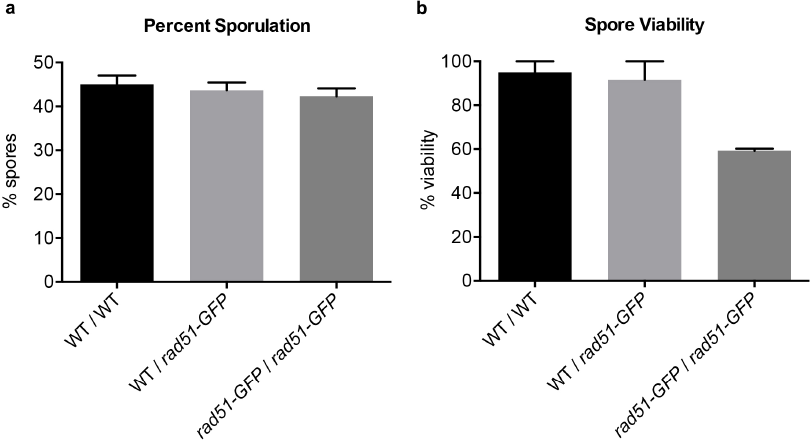
Rad51-GFP is competent in meiosis. **a)** Percent sporulated cells as determined by light microscopy in the indicated strain. **b)** Quantification of spore viability after tetrad dissection of sporulated cells. Error bars are representative of three biological replicates.

### Multiple DSBs form discrete Rad51-GFP foci

We extended our analysis to monitor the appearance of multiple DSBs, to determine whether multiple DSBs would appear as a single Rad51-GFP focus or as distinct Rad51-GFP foci. We inserted Rad51-GFP into strain YCSL004 carrying 3 HO cleavage sites, each on a different chromosome, as well as Rad52-RFP, and counted the number of foci 3 h after *GAL::HO* induction. We observed an average of 2 Rad51-GFP foci (Figures 4A, B, S2A). This distribution was unchanged in *lig4Δ* cells, in which repair by end-joining is blocked (35-37)(Figure S2B).

**Fig 4.**
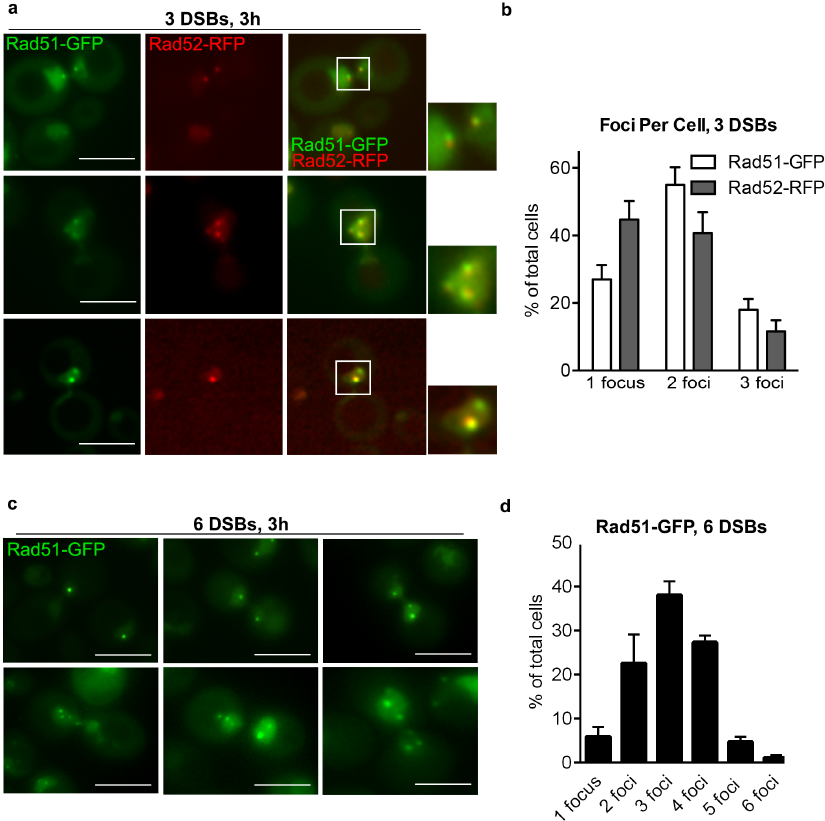
Rad51-GFP forms multiple foci in response to multiple DSBs. **a)**Representative images from strain DW106 expressing endogenous Rad51-GFP and Rad52-RFP from its endogenous promoter 3 h after HO induction. Images prepared as in Figure 1A **b)** Quantification of Rad51-GFP and Rad52-RFP foci in DW106 3 h after induction of HO **c)** Representative images from strain DW280 (6 HO cute sites) expressing endogenous Rad51-GFP 3 h after HO induction prepared as in (a) **d)** Quantification of foci per cell in DW280 as described in (c). Error bars represent the SD of three biological replicates of >150 cells per experiment.

In wild type cells, we noticed several instances of cells displaying a single Rad52-RFP focus but multiple Rad51-GFP foci (Figure 4Aiii and B). In these cells, the single Rad52-RFP focus was typically large and always colocalized with a single Rad51-GFP focus. Therefore, monitoring the number and locations of DSBs via Rad52 may not serve as a realistic reflection of the actual DSB state.

We also examined Rad51-GFP foci in strains with 6 HO cleavage sites (38). We observed a range of the number of foci per cell, averaging 3.2 foci, 6 h after HO induction (Figures 4C, D, and S2C).

### Live cell detection of DSBs with Ddc2-GFP

In strains with multiple DSBs, only a small proportion display a single Rad51-GFP focus, raising the question whether DSBs are usually recruited into a repair center. However, whether the distribution of foci depends on Rad52 is impossible to test using Rad51-GFP, as Rad51’s recruitment is completely dependent on Rad52 (28, 39). To directly ask whether Rad52 recruits multiple DSBs to the same location in the nucleus, we monitored DSB dynamics using a live cell marker that is independent of Rad52.

One such candidate is the DNA damage checkpoint protein Ddc2. Ddc2 localizes to a broken DNA end, either directly or by binding to RPA (22, 23, 40) and previous studies have shown strong localization of Ddc2-GFP at DSB sites (41-43). We carried out an analysis similar to that described above for Rad51-GFP, using a derivative of strain YSCL004 with 3 HO-induced DSBs but carrying an insertion of GFP at the C-terminus of the chromosomal copy of Ddc2 (strain VE290). Again, we observed cells with 1, 2, or 3 foci with an average of 2 foci per cell (Figures 5A, B, and S3A). We repeated this analysis in a *rad52Δ* derivative and found no difference in the distribution of Ddc2-GFP foci (Figures 5B and S3B). Hence, Ddc2 foci are not dependent on Rad52. However, in *rad52Δ,* we noticed a small percentage of cells that had more than 3 foci (Figure 5B, S3B, Movies S1-S3). In 5.4% of *rad52Δ*, cells, 4 and sometimes 5 foci were evident (Figure 5B), suggesting that Rad52 might be partially responsible for holding together the ends of DSBs.

**Fig 5.**
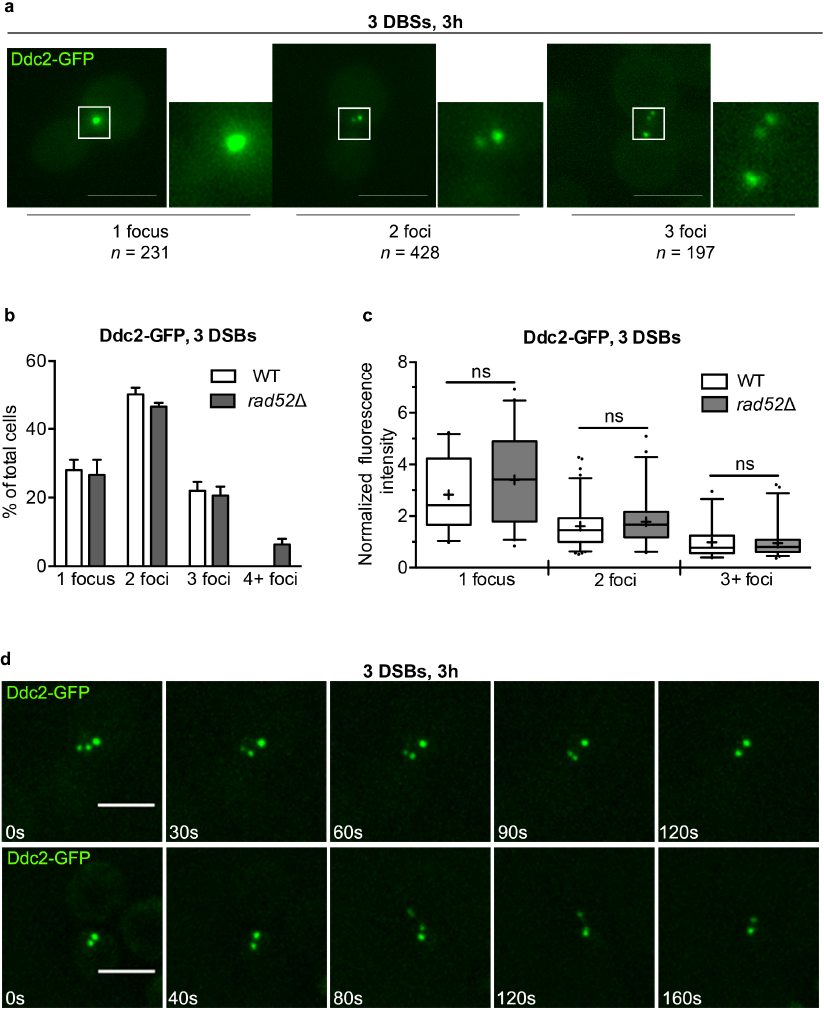
Analysis of Ddc2-GFP focus dynamics after 3 DSBs. **a)** Representative images of *1, 2,* or 3 foci in strain VE290 expressing endogenous Ddc2-GFP 3 h after HO induction. *n* = total number of cells displaying the indicated number of foci from three biological replicates. Images prepared as in Figure 1A. **b)** Quantification of Ddc2-GFP foci in strain VE290 (WT) and DW546 (*rad52Δ*) 3 h after HO induction. Error bars represent the SD of three biological replicates totaling 856 (WT) and 592 *(rad52* Δ) cells. **c)** Background-subtracted fluorescence intensities of individual foci in strains VE290 (WT) and DW546 (*rad52Δ*) 3 h after HO induction as described in B. Box plots prepared as in Figure 1E. **d)** Time lapse images of Ddc2-GFP in strain VE290 suffering 3 DSBs 3 h after HO induction. Time after first image displayed in seconds below.

Even in the absence of Rad52, we observed ~25% of cells displaying a single focus. It is possible that in single-focus cells, HO had not efficiently cut all three sites. To address this possibility, we took advantage of the high signal specificity and low nuclear background signal in cells expressing Ddc2-GFP to determine the fluorescent signal intensity of individual foci. It is evident that the intensity of the single focus was much greater than the average intensities of each focus in cells displaying 2 foci or 3 foci. Indeed, the signal intensity of 1 focus is equal to the sum of the signal intensities of 3 foci (Figure 5C). Thus, cells with a single focus apparently have 3 DSBs that are indeed co-localized. These intensities were unchanged in *rad52Δ* (Figure 5C).

### DSB foci are dynamic

Chromosomal mobility and chromatin persistence length are radically altered after the induction of a DSB (44-46). We examined the stability of foci with 3 DSBs DSB by observing cells using over a ten minute period 3 h after HO induction, using spinning disk confocal microscopy. In 85% of cases, the number and general localization of foci in a given cell remained constant over 10 min (Movies S7-S10). However, in 15% of cells, we observed changes in the number of Ddc2-GFP foci. We found instances when the number of foci diminished, from three to two or from two to one, as well as a single focus splitting into two or three foci (Figure 5E, Movies S11 – S17). This behavior was unchanged in *rad52Δ,* with the exception of a few cells with >3 foci described above. We conclude that DSBs are dynamic and that Rad52 is not the mediator of a DSB repair center.

### Microtubules control DSB dynamics

Due to the dynamic nature of multiple DSBs, we sought to determine the molecular mechanism behind this motion. The spindle pole, microtubules, and the kinetochore have all been implicated in governing chromatin mobility in response to DNA damage (47, 48). Furthermore, DSBs have been shown to colocalize with spindle pole bodies preferentially loaded with the SUN protein Mps3 (12, 49). However, in our 3-DSB system expressing Ddc2-GFP and Mps3-mCherry, we find only 28% of cells exhibit Ddc2/Mps3 colocalization (Figure 6A). To test whether the action of microtubules was required for DSB dynamics in our system, we induced HO for 3 h in cells suffering 3 DSBs and expressing Ddc2-GFP. After 2 h, we added nocodazole for 1 h then monitored foci dynamics by live cell confocal microscopy (Figure 6B). The distribution of foci per cell was drastically shifted towards many more single focus cells (Figure 6C). Thus, DSB dynamics are driven by microtubules and in the absence of microtubules multiple DSBs colocalize.

**Fig 6.**
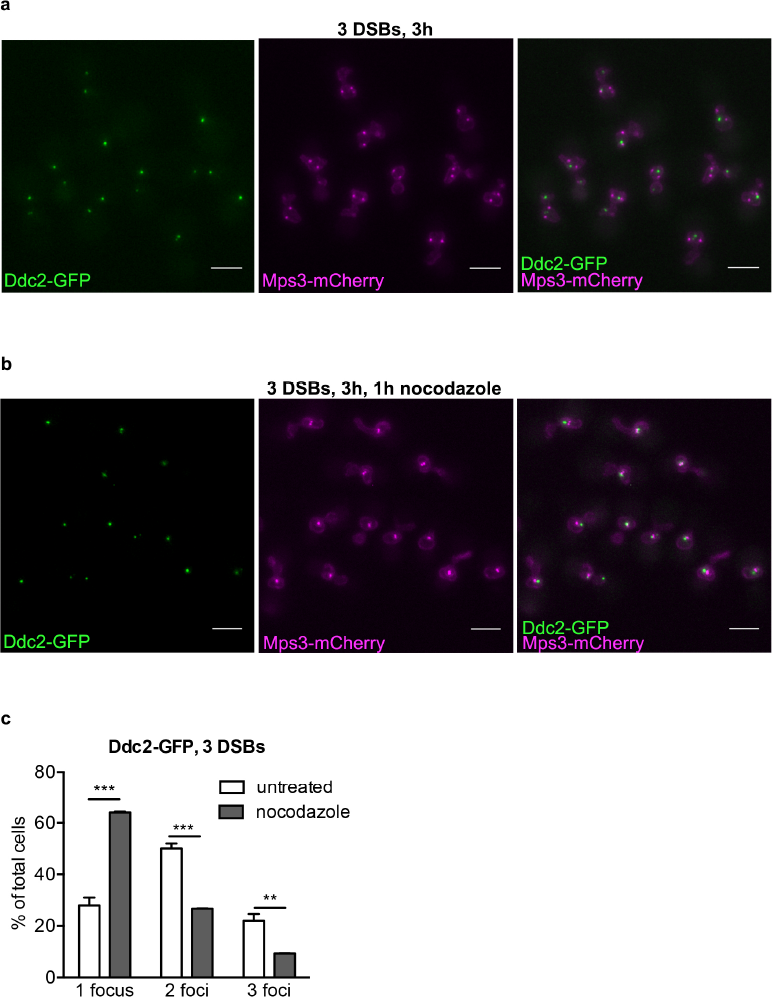
Microtubule dependent Ddc2 foci dynamics. **a)** Strain DW547 expressing Ddc2-GFP and Mps3-mCherry 3 h after HO induction. Images prepared as in Figure 1A. **b)** Similar to (a) with the addition of 15 µg/ml nocodazole after 2 h of HO induction. c) Quantification of Ddc2-GFP foci from cells in (b). Error bars represent the SD of three biological replicates of >150 cells per experiment. Student’s t test: *** p ≤ 0.001, ** p ≤ 0.01.

## Discussion

DSB repair must be coordinated in space and time in order to faithfully repair lesions to the genome. The role of many proteins involved in DSB repair has been elucidated through in vitro and in vivo biochemistry, but the lack of suitable live cells markers has provided a barrier to studying DSB repair in real-time. Here, we report DSB dynamics in single-and multiple-break conditions using two different fluorescently tagged proteins that carry out different functions in response to DNA damage; the recombinase Rad51-GFP and the checkpoint-related protein Ddc2-GFP. In both cases, multiple DSBs resulted in multiple fluorescent foci.

Using Rad51-GFP or Ddc2-GFP in our 3 DSB system, the majority of cells exhibit two or three foci. Rad51-GFP foci often colocalize with Rad52-RFP, but we see many instances with more GFP foci than RFP foci. Previous studies have looked specifically at the role of Rad52 in organizing a “repair center” yeast (15, 50). Our data suggests that monitoring Rad52 focus formation may underestimate the number of DSBs throughout the nucleus. This difference may in part reflect the temporal recruitment of DSB repair proteins to the site of DSBs such that the continued presence of Rad52 at a DSB may not be necessary once a Rad51 filament has been established.

While our Rad51-GFP construct is not able, by itself, to repair DSBs by homologous recombination, it is not dominant negative and supports recombination in meiosis. Biochemical work on a human isoform of Rad51 fused to GFP determined that the fluorescent tag prevented Rad51 from engaging in the pairing of homologous sequences by inhibiting Rad51’s secondary DNA binding (20). We envision the same to be true of our Rad51-GFP construct because our ChIP experiments and microscopy suggest that Rad51-GFP can efficiently bind to ssDNA and form a filament, its first step in homologous recombination. However, when Rad51-GFP is the sole copy of Rad51 in cells, DSB repair by homologous recombination is incomplete, presumably at the strand exchange step.

Rad51-GFP’s defect in ectopic gene conversion is suppressed by addition of a single second copy of wild type Rad51 expressed from its endogenous promoter. Likewise, the MMS sensitivity conferred by Rad51-GFP was also rescued by expression of wild type Rad51. However, it is not that wild type Rad51 simply excludes Rad51-GFP from binding ssDNA, since Rad51-GFP readily forms a focus in the presence of wild type Rad51. This is similar to a similar construct reported in fission yeast (21). Together with previous reports, data suggest either that a functional Rad51 filament does not require every Rad51 molecule to be functional or that subunit-subunit interactions between wild type and GFP-tagged Rad51 corrects the defect. The exact stoichiometry for a functional filament cannot be determined from these experiments, but from previous work done by our lab (51) we speculate that there need to be at least two to three functional Rad51 molecules in tandem to facilitate minimal Rad51-mediated strand exchange.

To directly test whether Rad52 recruits multiple DSBs into a common locus, we used Ddc2-GFP, which forms foci independent of Rad52. In our 3-DSB strain, we see an average of 2 Ddc2-GFP foci per cell, but still about 25% of cells display a single focus, as with Rad51-GFP. However, this distribution remains unchanged in a *rad52Δ* derivative. Therefore, we conclude that Rad52 is not required for organizing multiple DSBs into one specific nuclear location. However, Rad52 appears to be partially responsible for tethering the ends of the DSB together, as we see a small but significant population of cells with greater than 3 Ddc2-GFP foci in *rad52Δ.* Both the Ku complex and Mre11 have been implicated in DBS end tethering previously (1, 53). but this study is the first to suggest that Rad52 is also a key player in end tethering.

That DSB foci are dynamic also supports our model that DSBs do not generally form a repair center. Increased chromatin motion in response to a DSB is believed to aid in DNA repair through facilitating in homology search throughout the genome (44-46) but the precise mechanism for this motion is relatively unclear. A recent study from the Durocher lab demonstrated that the DNA damage checkpoint kinase cascade targets the kinetochore-associated protein Cep3 and this phosphorylation increases chromatin movement through activation of the spindle assembly checkpoint (47). Similarly, increased chromatin movement in response to a single DSB has been shown to be microtubule dependent (48). In our system, functional microtubules promote DSB dynamics; in their absence, DSBs tend to coalesce. However, given that the foci distribution in three HO break cells are not altered in *rad52Δ,* the exact mechanism behind this coalescence remains to be determined.

## Materials and Methods

### Strain and Plasmid Construction

Standard yeast genome manipulation procedures were used for all strain constructions (54). Linear DNA and plasmids were introduced by the standard lithium acetate transformation procedure (55). To C-terminally tag Rad51 and Ddc2 with GFP, PCR primers were used to amplify the GFP fragment from pFA6a-GFP(S65T) and the *TRP1* or KAN selectable marker in the Longtine collection (56) and introduced to the appropriate parent strain by lithium acetate transformation. Strain genotypes are listed in Table S1. Primer sequences are listed in Table S2.

### Growth Conditions

To visualize the chromosomally integrated fluorescent tags (Rad51-GFP and Ddc2-GFP) after DNA damage, cells from a single colony were grown overnight in 5ml YEP + 4% lactic acid (YPLac). Cells were diluted to OD600 = 0.2 and grown for 4 h in 5 ml of fresh YPLac before addition of galactose to a final concentration of 2% to induce *GAL::HO* expression. For experiments that visualized Rad52-RFP, the same growth procedure except that cells were grown in SD-leucine media supplemented with 2% raffinose.

### Plating Assays and Viability

The efficiency of DSB repair by homologous recombination was determined as described previously for strain YJK17 (32). Briefly, cells of the appropriate strain were selected from a single colony on YPD plates and grown overnight in 5 ml of YPLac. Cells were diluted to OD600 = 0.2 and allowed to grow until OD600 = 0.5 – 1.0. Approximately 100 cells from each culture were then plated on YPGal (2% v/v) and YPD in triplicate and incubated at 30 °C. Viability was calculated by dividing the number of colonies on YPGal by the number of colonies on YPD.

Adaptation assays in strain JKM179 were performed as previously described (24). Briefly, cells were grown in YPLac or SD-media supplemented with 2% raffinose overnight then individual unbudded (G1) cells were plated on YPGal and observed microscopically for 24 h to determine the percent that were arrested in the G2/M stage of the cell cycle.

Viability on MMS media was determined by as described by (57). Cells of the appropriate strain were selected from a single colony on YPD plates and grown overnight in 5 ml of selective media to near saturation. The following day, cultures were diluted to OD600 = 0.2 and left to grow at 30 °C for 3-5 doublings. Cells were then diluted in 200 µl sterile water to OD600 = 0.2 in a 96-well plate and subsequently 10-fold serially diluted six times. Cell dilutions were then plated on YPD,-leu, and -leu +0.002% MMS plates and left to grow at 30 °C for three days.

### Image Acquisition and Analysis

Prior to imaging, cells were washed twice in imaging media SC supplemented with 2% galactose or 2% raffinose and mounted on a glass depression slide coated with agarose supplemented with all amino acids. GFP and RFP signals were visualized with Zeiss AxioObserver spinning disk microscope, 63x objective set to acquire 10 z-stack images spaced at 0.4 µm. Z-stacks were imported into Fiji and max-projected to acquire a single image sum of all slices. Foci were counted by adjusting the image color threshold to the average nuclear signal intensity for a given image and counting spherical regions that gave pixel intensity above the threshold. For GFP and RFP colocalization analysis, max projected z-stacks were merged in Fiji and analyzed for overlapping foci. Rad51-GFP nuclear intensities were quantified by measuring the integrated intensities of concentric nuclear circles from max projecting z-stack images and subtracting from this value the average background fluorescent intensities. Ddc2-GFP spot intensities were determined in a similar fashion.

### Chromatin Immunoprecipitation

Chromatin immunoprecipitation (ChIP) was carried out as described in (58). In brief, cells were harvested from log-phase population. 45 ml of culture were fixed and crosslinked with 1% formaldehyde for 10 minutes after which 2.5 ml of 2.5 M glycine was added for 5 minutes to quench the reaction. Cells were pelleted and washed 3 times with 4°C TBS. Cell wall was disrupted by 1 h bead beating in lysis buffer, after which cells sonicated for 2 minutes. Debris was then pelleted and discarded, and equal volume of lysate was immunoprecipitated using α-ScRad51 antibody for 1 hour in 4°C, followed by addition of protein-A agarose beads for 1 h at 4°C. The immunoprecipitate was then salt washed 5 times, and crosslinking was reversed at 65°C overnight followed by proteinase-K addition for 2 h. Protein and nucleic acids where separated by phenol extraction. Chromatin association with Rad51 was assessed by qPCR. More detailed protocols and recipes are available upon request. α-ScRad51 antibodies were a generous gift of A. Shinohara (University of Osaka, Osaka, Japan) and from Douglas Bishop (University of Chicago, Chicago, Il).

### DSB Repair Analysis by qPCR

Monitoring repair kinetics by qPCR was performed as described previously (59). Single colonies were inoculated in 5ml of-leu dropout media with 2% dextrose and grown overnight at 30C. Overnight cultures were then diluted into 600ml of YPLac and grown into log phase. DSBs were induced by adding 20% galactose to a final concentration of 2%. To track the dynamics of DSB repair 50ml aliquots of each culture was collected every hour over 9 h. DNA was isolated using a MasterPure™ Yeast DNA Purification Kit (Epicentre cat. MPY80200). The repair product *MATa-inc* was amplified using primers MATp13 and MATYp4 with a SYBR Green Master Mix using a Qiagen Rotor-Gene Q real-time PCR machine. To quantify the relative amount of *MATa-inc* in each sample, Slx4p was used as a reference gene and was amplified using primers NS047-Slx4p7 and Slx4p1. Primer sequences are shown in Table S2.

## Supplementary Figure Legends

**Table S1.**
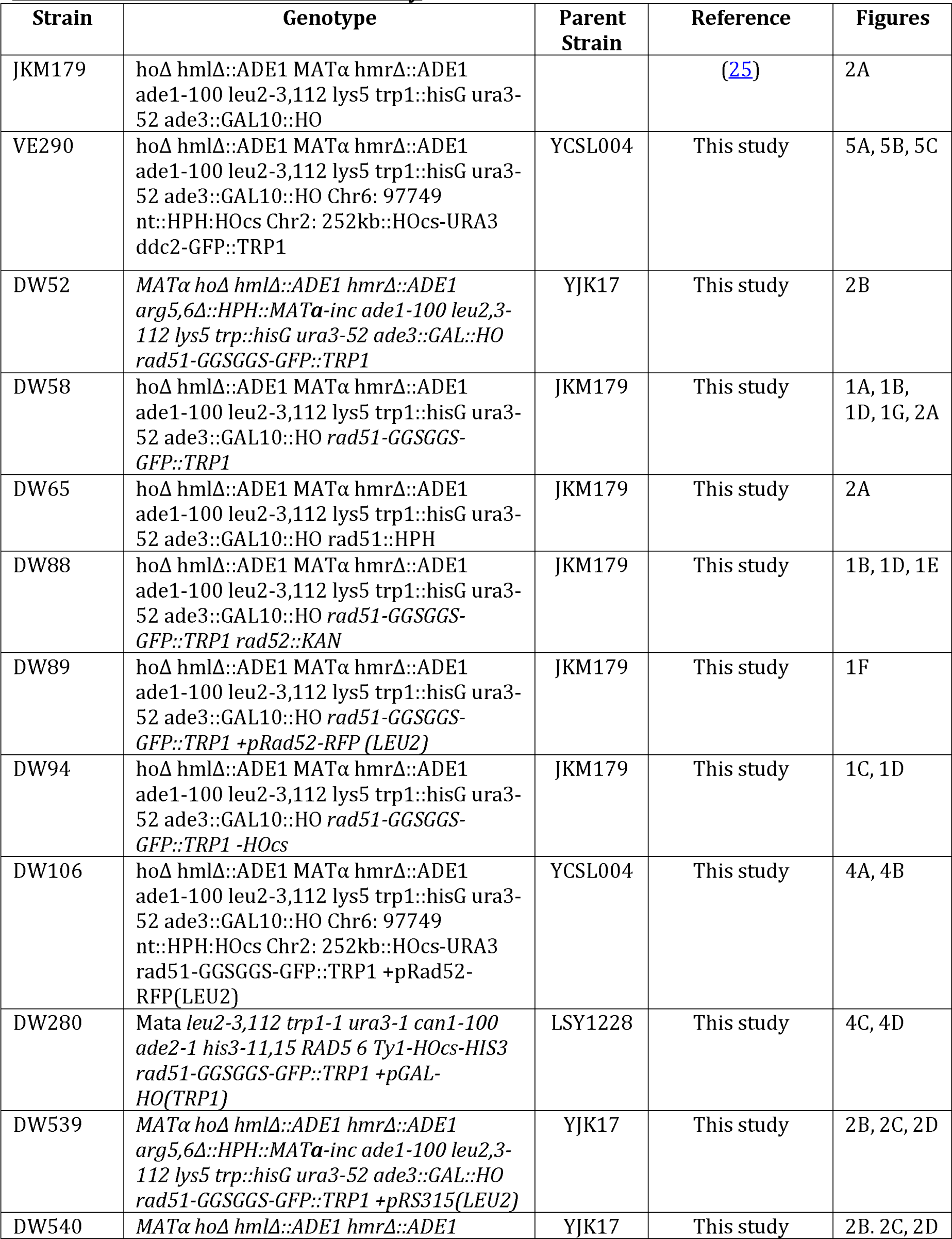
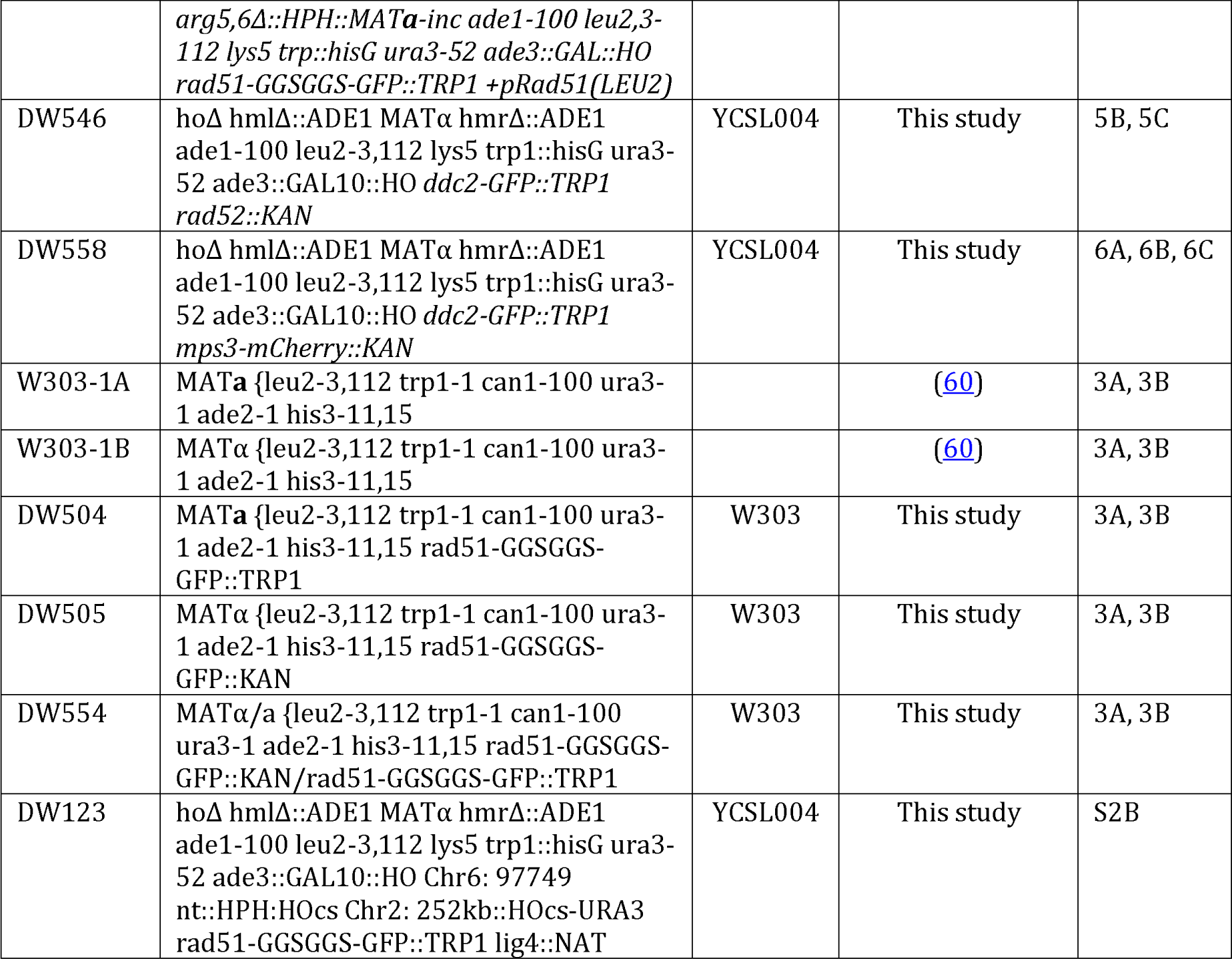
Strains used in this study

**Table S2.**
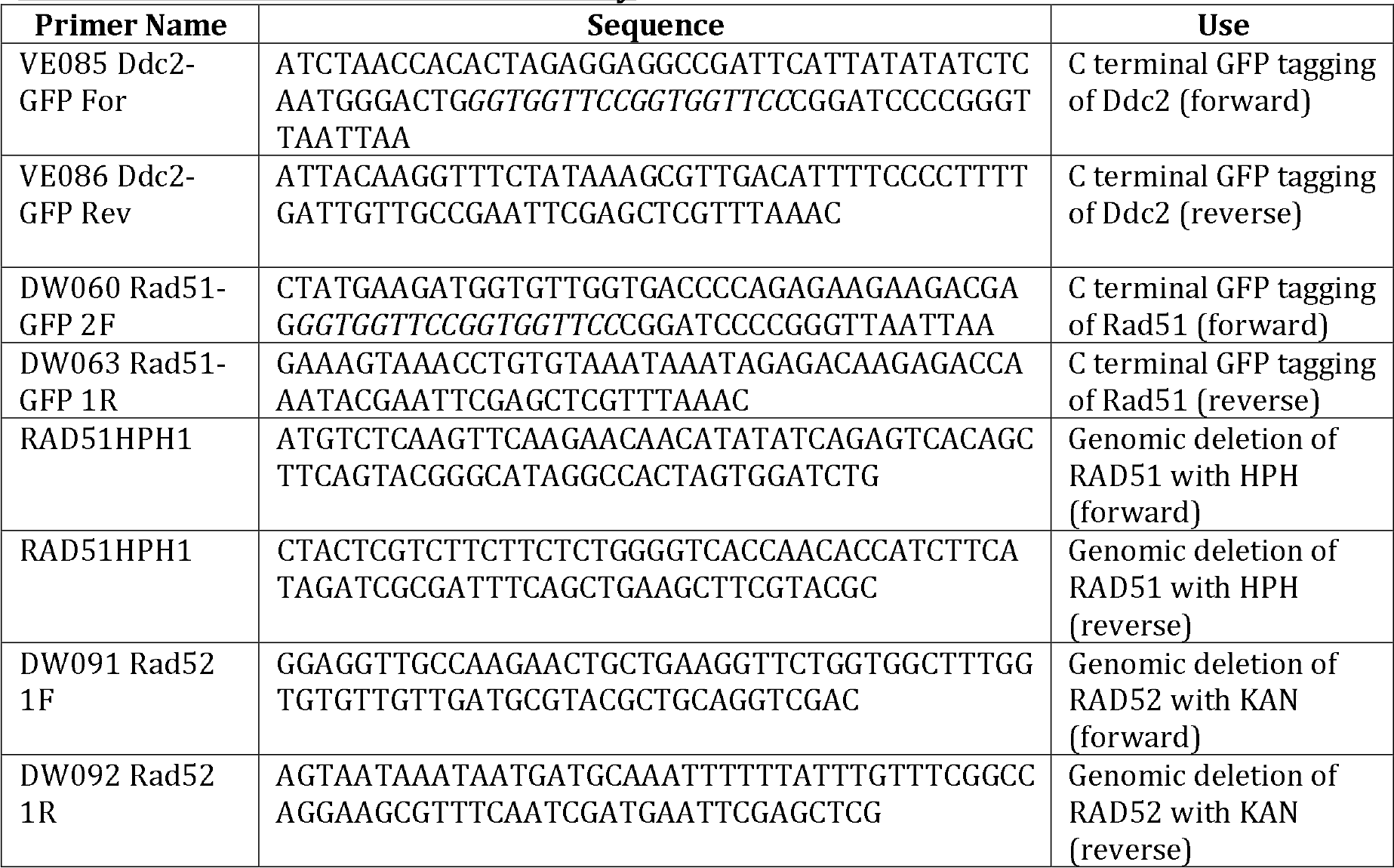
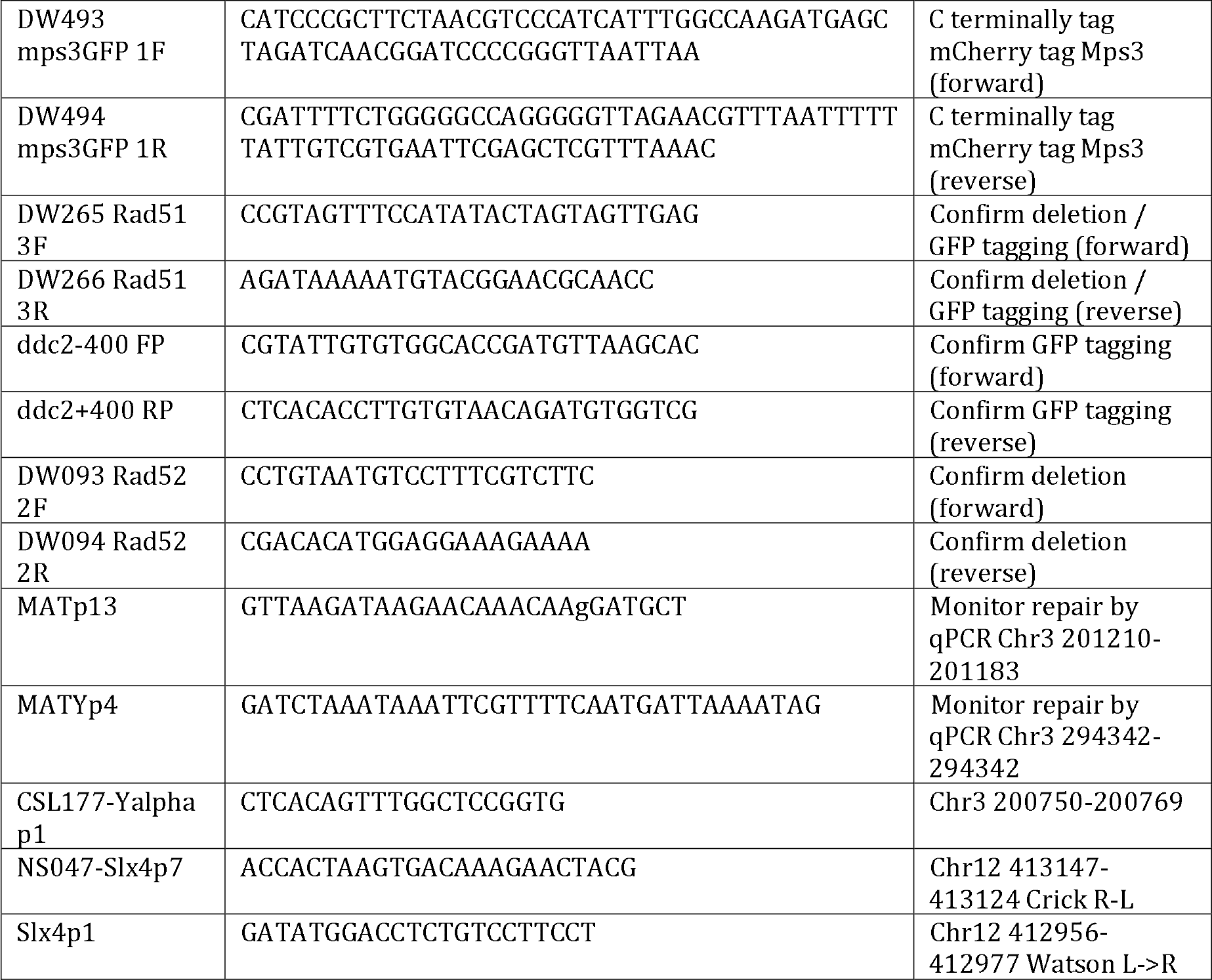
Primers used in this study

**Figure S1: Rad51-GFP localization._a)** Representative full field image of strain DW58 expressing endogenous Rad51-GFP 6 h after HO induction. **b)** Representative full field image of strain DW88 (*rad52Δ*) expressing Rad51-GFP 6 h after HO induction. **c)** Representative images of strain DW94 (no HO cut site) expressing Rad51-GFP 6 h after HO induction. **d)** Representative full field image from strain DW89 expressing endogenous Rad51-GFP and Rad52-RFP from its endogenous promoter on a low copy plasmid 3 h after HO induction. Maximum projection of 12 z-stack images every 0.4 µm. Scale bar = 5 um

**Figure S2: Rad51-GFP localization in multi-break strains** a)_Representative full field images of strain DW106 expressing Rad51-GFP and Rad52-RFP 3 h after HO induction. b) Quantification of Rad51-GFP foci in strain DW123 (*lig4Δ*) **c)** Representative full field images of strain DW280 expressing Rad51-GFP 3 h after HO induction. Maximum projection of 12 z-stack images every 0.4 um. Scale bar = 5 µm

**Figure S3: Ddc2-GFP localization in multi-break strains_a)** Representative full field image of strain VE290 expressing Ddc2-GFP 3 h after HO induction. **b)** Representative full field image of strain DW546 *[rad52*Δ) expressing Ddc2-GFP 3 h after HO induction. Maximum projection of 12 z-stack images every 0.4 μm. Scale bar = 5 μm

**Movie S4 – S6** Ddc2-GFP in 3 DSB strain DW546 (*rad52Δ*) 3 h after HO induction. Scale bar = 5 µm.

**Movie S7 – S17** Ddc2-GFP in 3 DSB strain VE290 3 h after HO induction. Scale bar = 5 µm.

